# Spatial proximity and dyadic social relationships affect ungulate behavioral synchrony

**DOI:** 10.1101/2025.08.06.668986

**Authors:** George M. W. Hodgson, Kate J. Flay, Tania A. Perroux, Alan G. McElligott

**Affiliations:** Department of Infectious Diseases and Public Health, Jockey Club College of Veterinary Medicine and Life Sciences, City University of Hong Kong, Hong Kong SAR, China; Centre for Animal Health and Welfare, Jockey Club College of Veterinary Medicine and Life Sciences, City University of Hong Kong, Hong Kong SAR, China; Department of Veterinary Clinical Sciences, Jockey Club College of Veterinary Medicine and Life Sciences, City University of Hong Kong, Hong Kong SAR, China

**Keywords:** “*Bos taurus*”, “cohesion”, “dominance”, “foraging”, “group coordination”, “ruminant”, “sexual segregation”

## Abstract

Collective group decisions are important for the survival and reproduction of social mammals, with inter-individual interactions often driving group-level emergent behavior. Activity synchronization is an important collective behavior, with differences in nutritional requirements leading to foraging asynchrony. Individual variation between animals (such as sex or social relationships) are predicted to affect ungulate synchronization and spatial proximity, with between-sex differences consequently influencing sexual segregation evolution in ungulates. Although investigated independently, the relative roles of sex, sociality and proximity in synchronization are rarely investigated concurrently, especially in regards to affiliative relationships. Asynchrony influences fission-fusion dynamics and social segregation, but little is known how short-term changes in synchrony affects fission. Using a mixed-sex group of feral cattle (*Bos taurus*), we evaluated the supporting evidence for several predictions arising from the current understanding of synchronization in ungulates. We investigated if sex and social relationships (dominance and affiliation) affected foraging, behavioral synchrony and proximity. We also investigated whether group synchrony affected short-term changes in group size (fission events). We found that same-sex dyads were more likely to be synchronized than mixed-sex dyads, but differences in dominance and affiliation did not affect dyadic synchrony. Focal animals were more synchronized with closest neighbors than with another randomly selected conspecific. Reduction in group size was more likely when group synchrony was lower, highlighting the importance of asynchrony in temporary movement decisions. Inter-individual differences can explain variation in collective behavior, with synchronization being biased towards certain individuals by favoring animals in close spatial proximity and those of the same-sex.

**LAY SUMMARY:** In ungulates, differences in energetic requirements lead to variation in activity, resulting in social and sexual segregation. However, sex, social relationships and spatial proximity are rarely investigated concurrently in relation to synchrony. We investigated synchronization and fission in a feral ungulate relative to individual differences in social relationship and proximity. Sex and proximity affected synchrony and fission events were more likely when synchrony was lower, highlighting the underlying processes in the evolution of sexual segregation.

## INTRODUCTION

Living in cohesive social groups provides considerable functional benefits to individuals, with coordinated movement maintained by collective behaviors (Ward & Webster, 2016). These collective behaviors help to transfer information and maintain group cohesion, with individual decision-making processes and inter-individual interactions often driving group-level emergent properties (e.g., group synchronization and fission-fusion dynamics; Ioannou and Laskowski 2023). Factors such as spatial proximity consequently affect the regulation of collective movement, coordination and the spread of information throughout a group, but there is relatively little information on how social relationships can mediate this effect (Amichay et al., 2024). Animals often have to differentiate between activities with varying costs and benefits and make behavioral decisions based on these trade-offs. Individuals may face consensus costs if their interests do not align with the majority of their peers (e.g., an individual is still hungry but the group has finished foraging) and they may be forced to forfeit the advantages of social living if they deviate from the group consensus (e.g., increasing predation risk if the individual remains foraging alone; King et al. 2008; Kerth 2010; Sueur and Deneubourg 2011). Understanding these trade-offs and the underlying processes that affect collective movement therefore allows us to comprehend different scales of behavior and decision-making in group-living social animals.

Collective group movement can arise from behavioral synchronization, occurring when two or more individuals in the same location conduct the same action simultaneously (Asher & Collins, 2012; Duranton & Gaunet, 2016). Synchronization of different behaviors is widespread across taxa and activity, from primates synchronizing their steps when walking together (Schweinfurth et al., 2022) to fish synchronizing their migration (Hulthén et al., 2022). Behavior events can also be unintentionally contagious, with nonconscious behavioral mimicry influencing individuals to change their activity to become similar to that of social partners; this social contagion can thus lead to the synchronization of all individuals in a group (Lakin et al., 2003; McDougall & Ruckstuhl, 2018; Ostner et al., 2021). However, synchronization is rarely examined concurrently relative to the characteristics of nearby social partners and their physical proximity. By helping individuals stay in close proximity, synchrony in group movement can help reduce mortality risk, enhance vigilance and deter predators (Boinski, 1987; Pays et al., 2007; Prokopenko et al., 2024). Through enhancing foraging, reducing predation risk, and helping maintain affiliative bonds, synchrony has large functional benefits for individual survival and reproduction (Jackson et al., 2008; Tarr et al., 2016).

Differences in individual physiology (such as those related to sex) are predicted to affect the likelihood of synchrony and proximity between two animals but sex, synchrony and proximity have rarely been examined concurrently (Bowyer et al., 2020; Michelena et al., 2004; Šárová et al., 2007). Synchrony may be costly to individuals if it requires them to make a choice between staying synchronized and a non-synchronized action with a higher benefit to themselves, as not all animals benefit equally from performing the same behavior (Ruckstuhl, 1998). This trade-off is often greater when individuals are of different ages, sexes or reproductive status with different nutritional requirements; for example, sexually dimorphic ungulate males and females differ in rumen size and nutrient requirements (Bowyer et al., 2020). Males therefore eat lower quality forage but require a greater amount than females, leading to the sexes being unable to synchronize their activity (Barboza & Bowyer, 2000; Mooring et al., 2005). Conflict in decision-making and asynchrony between individuals of different characteristics can result in fluctuating group membership, fission-fusion dynamics, and eventually in segregation (Bonenfant et al., 2004; Mooring et al., 2005; Ruckstuhl & Neuhaus, 2002).

Individual variation in social phenotype (e.g., dominance rank and social bonds) is likely to affect animal movement, proximity and synchronization (Connor et al., 2006; Fele et al., 2025; Krueger et al., 2014). Dominance can affect foraging activity, with higher-ranked individuals monopolizing higher-quality foraging sites and forcing lower-ranked animals into areas of lower quality forage or higher predation risk (K. J. Schneider, 1984; Waite, 1987). Subordinate animals may experience socially mediated interference while foraging, occurring when a dominant individual reduces another animal’s energy intake and leading to subordinates having lower energy reserves (Rands et al., 2006). Social relationships (e.g., affiliation, familiarity and personality) affect the intensity of synchronization (Gygax et al., 2010; Maeda et al., 2021), the likelihood of proximity (Briard et al., 2015; Keshavarzi et al., 2023) and behavioral contagion (Valente et al., 2022), but it is often difficult to determine affiliative relationships in wild ungulates. Proximity itself also affects coordination and synchrony (Amichay et al., 2024), and synchronization can arise through social facilitation, with individuals synchronizing their activity with the animals closest to them (Evans et al., 2018; Hoyle et al., 2021; Rands et al., 2014). However, individuals may mediate their responses to nearest neighbors depending on their identity, and the extent of the impact of social status on synchrony remains unknown (McDougall & Ruckstuhl, 2018; Valente et al., 2022).

Differences in energetic requirements, activity budgets and synchronization are key factors affecting fission-fusion dynamics and sexual segregation in ungulates (Fig. 1: Conradt 1998; Ruckstuhl and Neuhaus 2002; Sueur et al. 2011). Fission (the temporary splitting of a group) is predicted when differences in physiology and energetic requirements between individuals become too great, and can result in single-sex groups synchronizing their activity together (Sueur et al., 2011; Sueur & Deneubourg, 2011). Individual consensus costs of synchronization are higher in mixed-sex groups (Bonenfant et al., 2004; Conradt, 1998), and heterogenous groups may therefore be prone to forming temporary subgroups in accordance with individual requirements (Conradt & Roper, 2000; Kerth, 2010). Fission events are more likely when behavior is asynchronous (Busia et al., 2022), but fission decisions are also affected by predation risk, quality of social relationships, habitat heterogeneity and food competition (Bond et al., 2019; Della Libera et al., 2023; Le Goff et al., 2024; Sueur et al., 2011). The role of synchrony in short-term fission-fusion events remains largely unexplored but may ultimately affect the evolution of sexual segregation in ungulates living in mixed-sex groups (Busia et al., 2022; Conradt & Roper, 2000).

**Fig. 1:**
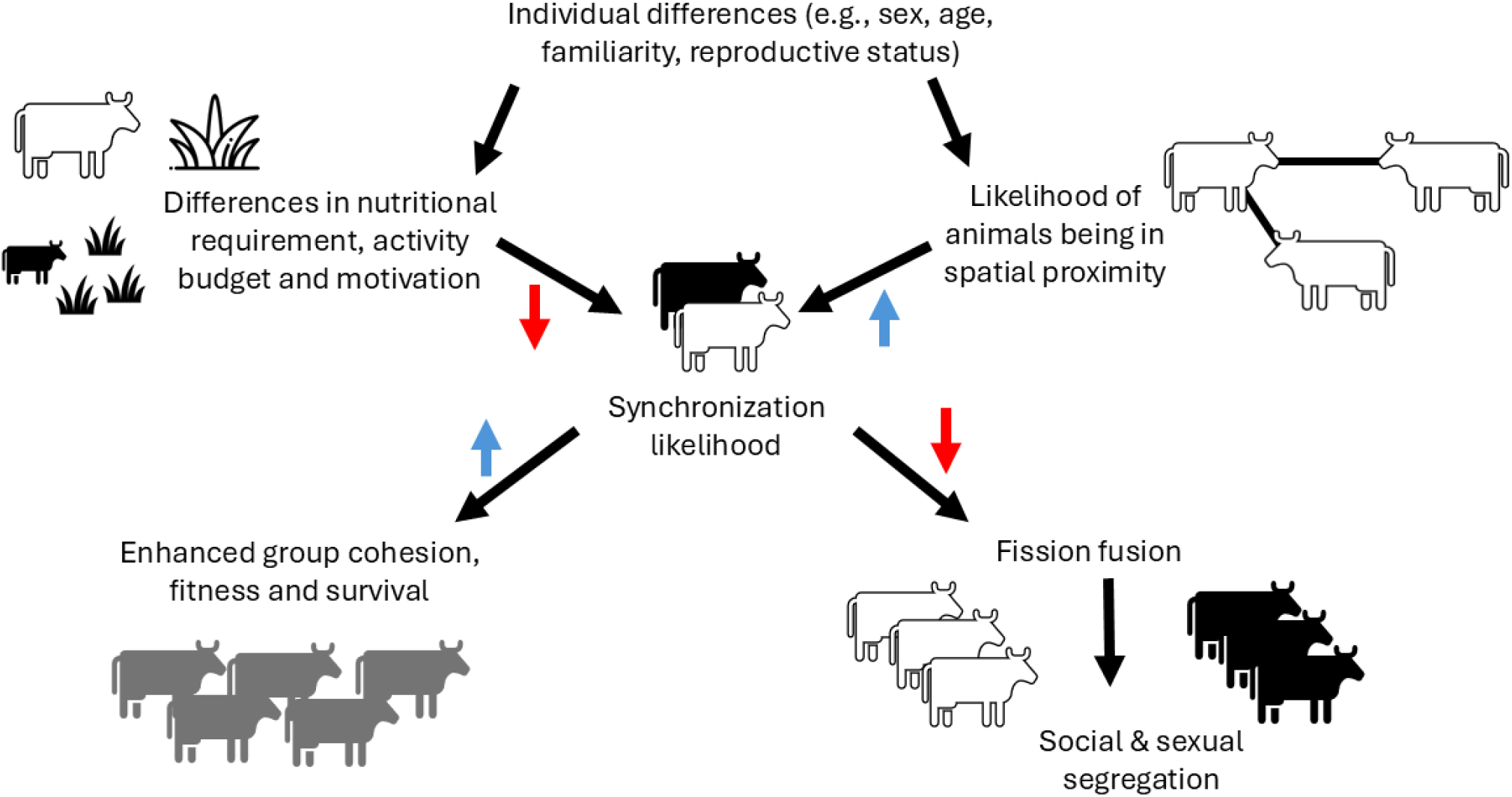
Individual differences between animals can affect differences in nutritional requirements and the likelihood of animals being in spatial proximity, but these factors are rarely addressed at the same time. These can affect synchronization likelihood, with lower synchronization leading to social segregation, and higher synchronization providing functional benefits (e.g., enhanced group cohesion).

Cattle (*Bos taurus*) are highly social, sexually dimorphic mammals, and engage in both affiliative and agonistic behavior. With clear dominance hierarchies and stable social relationships, cattle present an excellent opportunity to investigate how individual characteristics affect behavioral synchrony in a globally abundant ungulate (Bouissou et al., 2001; Polák & Frynta, 2010). In beef and dairy cattle, behavioral synchronization is related to time, proximity and similarity in body size, but not dominance status or reproductive state (Šárová et al., 2007; Stoye et al., 2012). With the majority of cattle research performed on cattle in farms, previous studies often examine synchronization from a productivity and animal welfare viewpoint. However, farm cattle social organization and space use is affected by husbandry, with food delivery times and movement restrictions affecting social interactions and food competition (Bouissou et al., 2001). This in turn can artificially synchronize activity, with farm cattle synchrony being affected by space availability (Færevik et al., 2008; Nielsen et al., 1997), group size (Schneider et al., 2020), and environment (Krohn et al., 1992; Tuomisto et al., 2019).

In Hong Kong, cattle were once used as draught animals until the decline of agriculture led to their release into surrounding areas during the 1950s - 1970s (AFCD, 2024; Barbato et al., 2020). Since then, animals are free-ranging in certain districts of Hong Kong, without routine farm husbandry and without any remaining large natural predators (Dudgeon & Corlett, 1994; Perroux et al., 2025a). Nowadays, free-ranging feral cattle live together for many years, with stable dominance hierarchies and variation in social bond strength (Hodgson et al., 2025a, 2025b). Although Hong Kong feral cattle are sexually dimorphic in body size (with males being larger than females), this dimorphism is moderate compared to other large ungulates, including most cattle breeds (Perroux et al., 2025b).

We examined the relationships between dyadic sociodemographic factors, spatial proximity and behavioral synchronization. We investigated several predictions arising from the current understanding of synchronization in ungulates, where synchrony and fission-fusion dynamics are both expected to impact the evolution of sexual segregation. Specifically, we first predicted (i) that sex and dominance rank would affect foraging activity, with males and higher-ranking animals spending less time foraging than females and lower-ranking animals (Ruckstuhl & Neuhaus, 2002). Second, we examined the role of spatial proximity in synchronization and predicted (ii) that spatial proximity would affect the likelihood of behavioral synchronization, with cattle synchronizing their activity with their nearest neighbors (Hoyle et al., 2021; Rands et al., 2014). Third, we examined the effects of sociodemographic factors on synchrony and proximity; we predicted (iii) that similarities in sex, affiliation and dominance would affect the likelihood of synchrony and close proximity between a dyad, with dyads composed of the same sex, of similar rank, or having previously exchanged affiliation being more likely to be synchronized and in close proximity (Connor et al., 2006; Conradt, 1998). Finally, we predicted (iv) that group synchrony would affect the likelihood of fission and reductions in group size, with animals being more likely to leave a group if the overall group activity was non-synchronized (Busia et al., 2022). Examining these factors concurrently allows a unique investigation into the driving forces of synchronization in ungulates.

## METHODS

### Ethical statement

Observational behavioural data were collected from a minimum distance of 15 – 20 m, adhering to the ethical guidelines outlined by the Association for the Study of Animal Behaviour to minimize disturbance to the animals (ASAB Ethical Committee & ABS Animal Care Committee, 2022). This work was approved by the Animal Research Ethics Sub-Committee of City University of Hong Kong (Internal Reference: A-0826).

### Study area and subjects

We collected observational data on a mixed-sex group (herd) of feral cattle, located in Kuk Po, Plover Cove, in the northeastern New Territories of Hong Kong. Cattle were observed throughout the study period in an area of approximately 70 ha (22°31’46"N 114°14’05"E), with surrounding areas consisting of brackish wetland, mangroves, small village houses, woodland and coastland. Several hiking trails run through the study site, with cattle habituated to human presence. There were 18 individuals observed at the study site over the study period (13 females, 5 males). This group was selected due to their known social relationships, high location fidelity, and because cattle did not receive regular organized provisioning with supplementary food; this group is also representative of the mixed-sex social structure and average herd size relative to other Hong Kong populations (Hodgson et al., 2025a; Perroux et al., 2025b). One male was observed in the group for a maximum of 9 minutes on one day only; this male and any observations during this period were removed from the dataset to avoid any potential bias towards rarely seen animals, resulting in 17 animals used in data analysis. Individual cattle age was unknown but only cattle over 1 year in age (identified by emerged horns and comparable size to known individuals) were observed in the group over the study period. Calves from multiple females have previously been observed in this group but no calves were present during the study period (Hodgson et al., 2025a). Hong Kong cattle are routinely sterilized for population management (AFCD, 2024), and although exact sterilization and reproductive states were unknown, the presence of calves in this group indicates breeding activity with multiple animals being non-sterilized. The mean observed group size during observations was 13.61, ranging from 8 to 17 animals. As Hong Kong cattle have a wide range of phenotypes (Barbato et al., 2020; Perroux et al., 2025b), cattle are individually identifiable and distinguishable via physical appearance (such as body size, horn shape, and coat color), with identities known from previous observations (Hodgson et al. 2025).

### Behavioral observations and data collection

We used instantaneous scan sampling to record data between 13 March 2023 and 24 March 2023 using mobile recording software ZooMonitor (Ross et al., 2022), with 128 sessions of 20 minutes duration (Altmann, 1974). Each 20-minute session consisted of 4 scans, sampling every 5 minutes with a total of 512 complete group scan samples. For each session, a focal animal was selected using a random generator from the list of animals which were present and had not been a focal in the previous scan. The average number of sessions per focal animal was 7.53 ± 1.17 (standard deviation, SD), ranging from 4 to 9 sessions. The behavior and identities of all individuals present were recorded for each scan, specifying the animal identity for the focal’s nearest neighbor one (NN1), nearest neighbor two (NN2), and nearest neighbor three (NN3) (Hoyle et al., 2021; Rands et al., 2014). We also recorded the proximity of each animal in the herd (within 1 body length [average body length of animals in this herd is approximately ∼ 1 m], within 3 body lengths [∼ 3 m], over 3 body lengths [over ∼ 3 m]) to the focal individual (Perroux et al., 2025b). Each individual was observed for 5 seconds to determine their active behavior, with mutually exclusive behaviors classified into four categories (lying, standing [resting, ruminating or vigilant while standing], walking or foraging [grazing and browsing, including drinking]). All individuals within view of the observers (with observers positioned at opposite ends of the group) and the other animals were recorded in the scan as part of the group. In addition to the 128 focal sessions used for data analysis, sessions where cattle were externally disturbed by human or dog activity were terminated early and discarded from analysis (13 sessions), as well as sessions missing the behavior of all group members present or without the identity of the focal animal’s nearest neighbors due to visual obstruction (7 sessions). Each session occurred between 09:00 and 17:00, with observations condensed into 10 days to minimize any impact from changes in group composition, temperature, weather, or reproductive state (Hoyle et al., 2021). Hong Kong weather in March is at the end of the dry season, with March 2023 having a mean monthly temperature of 21.3 degrees, total monthly rainfall of 70.3mm, and 76% mean relative humidity (Hong Kong Observatory, 2024).

### Data preparation

To examine how synchrony was affected by dyadic social relationships, we used existing social relationships as a measure of dyadic affiliation, and calculated the dominance rank of each individual within the group (Hodgson et al., 2025a). These supporting behavioral interaction data were collected in addition to the behavioral synchrony data during an independent previous observation period between 17 August 2022 and 22 May 2023, with 44 hours of all-occurrence sampling between 10:00 and 16:00 (Hodgson et al., 2025a). Although the synchrony data collection falls within this time period, no data were included in both observations. Supporting behavioural interaction data was collected by a familiar observer using mobile recording software ZooMonitor (Ross et al., 2022), recording allogrooming events (social grooming) and non-contact displacement behaviors (Hodgson et al., 2025a). Group size ranged from 7 to 20 animals, with all animals recorded in the synchrony data also being observed in the supporting behavioural interaction data. Allogrooming was defined as repeated licking by one animal on another animal’s body, and distinct bouts separated by a break of 10 seconds or longer (Hodgson et al., 2024; Laister et al., 2011). Dyadic affiliation was therefore defined as a binary measure of whether two animals had previously exchanged allogrooming. Dominance interactions were also recorded over this period, with non-contact displacement behaviors defined as the approach of one animal (the ‘winner’) causing another animal (the ‘loser’) to withdraw from their original position, and take at least three steps away from the performer (Christensen et al., 2002; Hodgson et al., 2024). These directed behaviors were transformed into a winner-loser matrix, and we used the randomized Elo-ranking methodology with 10000 randomizations via R package ‘aniDom’ in R to calculate dominance scores for each individual (Farine & Sánchez-Tójar, 2021; Sánchez-Tójar et al., 2018). We then converted the dominance scores into ordinal dominance ranks from 1 to 17, with 1 indicating the highest-ranked individual in the group.

### Statistical analysis

Data calculations and analyses were performed using R version 4.4.3 (R Core Team, 2024), with data and code accessible via an online OSF repository (Dataset, 2025). Synchrony measures are sensitive to both group size and the number of activity categories, and we were unable to fully calculate the Fleiss Kappa coefficient due to fluctuating group membership between scans (Asher & Collins, 2012; Stoyan et al., 2018). Instead, we partially calculated a Fleiss Kappa coefficient of agreement for 33 scans in which all 17 animals were present, using function ‘*kappam.fleiss’* from package ‘irr’ (Gamer et al., 2005). In addition, for all scan observations we calculated group synchrony as the maximum percentage of animals simultaneously performing the same behavior per scan, and we calculated the mean degree of group synchrony across all observations. We also calculated the proportion of observations which were equal to or higher than thresholds of 70%, 90% and 100%, where all cattle were in agreement. These thresholds were chosen due to cattle being at least 70% synchronous on the majority of occasions in previous research (Stoye et al., 2012). All models were tested to ensure they fit model assumptions using packages ‘DHARMa’ and ‘performance’ (Hartig & Lohse, 2022; Lüdecke et al., 2024). *P* values were obtained through function ‘*drop1’* from package ‘stats’, with single-term deletions for all generalized linear models (GLMs) and generalized linear mixed-effects models (GLMMs), and obtained using Satterthwaite’s single-term deletions for linear mixed-effects models (LMMs).

#### Effect of sex and dominance on foraging (i)

To test whether sex and dominance affected foraging activity, we calculated the proportion of scans that each animal spent foraging; this was defined as the number of times that an animal was observed foraging divided by the individual’s total number of observations. We tested the effect of sex and dominance rank on the proportion of time spent foraging using a GLM from R package ‘glmmTMB’ with a Gaussian distribution and identity link (Brooks et al., 2023).

#### Effect of spatial proximity on synchrony (ii)

We investigated whether focal animals were more synchronized with their nearest neighbors or with another random animal in the group (random conspecific). The conspecific was selected by randomly sampling an individual present during each scan that was not a near neighbor (nearest neighbor 1, 2, or 3); this sampling was performed post-data collection using R to avoid any visual biases that may arise from random selection of animals on field. We then calculated the proportion of scans in each session (*n*/4) that each type of animal (neighbor 1, neighbor 2, neighbor 3, random) was synchronized with the focal animal (Hoyle et al., 2021; Rands et al., 2014). Synchrony therefore ranged from 0.00 (activity was never matched with focal’s activity) to 1.00 (activity was always matched with focal’s activity). We used a GLMM with function ‘*glmer’* from ‘lme4’, using the proportion of synchronized scans as a dependent variable weighted by the number of scans (number of synchronized scans / 4), and included the type of animal as an independent variable, with a logit link. Hour of observation and focal animal identity were fitted as random effects.

#### Effect of sex, dominance and affiliation on synchronization and proximity (iii)

To investigate whether synchrony between the focal animal and their nearest neighbor was affected by their sex, dyadic affiliation and dominance rank difference, we used each scan with a focal and their nearest neighbor (512 events). We classed each dyad as either same-sex (female-female or male-male) or different-sex (male-female). We assigned each dyad a binary measure of affiliation (whether the pair had previously exchanged allogrooming (1) or they had not (0)), and calculated the absolute dominance rank difference between their two ordinal ranks. We used a GLMM (function ‘*glmmTMB*’ from R package ‘glmmTMB’) with a binomial family and logit link, testing whether the predictor variables of dyadic sex combination (same-sex or different sex), previous affiliation (1 or 0), and dominance rank difference affected the binomial response of whether the two animals were synchronized (1) or not (0). The distance of the nearest neighbor from the focal (within 1 body length, within 3 body lengths, over 3 body lengths), the identities of both animals, and the hour of observation were fitted as random effects.

To test whether overall synchrony between two animals was related to their sex, dyadic affiliation and dominance rank differences, we recorded all possible dyadic pairs of animals (136 dyads). We then calculated dyadic synchrony as the proportion of scans where both animals were present and performing the same behavior (Hauschildt & Gerken, 2015; Šárová et al., 2007). We used a LMM with ‘*lmer’* from R package ‘lmerTest’ (Kuznetsova et al., 2020), to test whether dyadic sex combination (same-sex or different sex), previous affiliation (1 or 0), and absolute dominance rank difference affected the proportion of scans synchronized, with both cattle identities entered as random factors (Šárová et al., 2007). For comparison, we calculated the dyadic expected degree of synchronization which would occur by two animals (A and B) behaving independently (Šárová et al., 2007). This was calculated as:

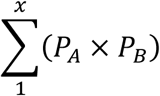

where *x* was the number of behavioral categories (4; lying, standing, foraging, walking), and *P* was the proportion of scans that A and B spent active in *x* behavioral category. We used a paired t-test to test whether dyads differed in their expected synchrony versus their observed synchrony, via function ‘*t.test*’ in package ‘stats’.

To test whether sex, dominance and affiliation affected the likelihood of proximity between each dyad, we used body length distance between the focal animal of each session and all other animals. We calculated the likelihood of a dyad being in close proximity as the number of times that both animals were recorded as within three body lengths of each other, divided by the total number of times that they were both present and one was a focal animal. We then ran a LMM to test whether the fixed variables of dyadic sex combination (same-sex or different sex), previous affiliation (1 or 0), and absolute dominance rank difference affected the likelihood of being in close proximity, with both animal’s identities fitted as random factors.

#### Effect of synchrony on likelihood of group fission (iv)

To test whether synchronization affected the likelihood of individuals leaving the group, we calculated whether group size changed between scans. We discarded all first scans in sessions due to having no previous comparable group membership size for sessions that started on different days or sessions that were not consecutive. This resulted in three scans per session in which we are able to calculate group size changes (384 scans). Net group size change was classified as a binary factor, with each scan being either 0 (no group size change from previous scan) or 1 (group size has net reduced since previous scan, with one or more animals leaving the group). There were 22 occurrences of the same number of animals leaving the group and rejoining the group at the same time; as these could not be classified as solely fission or fusion, these were removed from the dataset and resulted in 362 scans used in the final analysis. We calculated group synchrony per scan as the maximum percentage of animals simultaneously performing the same behavior (number of animals performing the same behavior divided by the scan group size). We also calculated the sex ratio of each scan by the number of males present divided by the number of females present. We ran a binomial GLMM with logit link from ‘lme4’, testing if the fixed factors of maximum group synchrony and sex ratio affected group size change from the previous scan, with group size and session identity as random factors.

## RESULTS

### Synchronization and sex differences in foraging (i)

Feral cattle were highly synchronized, with an average group synchrony degree of 79.89% (range 28.57% to 100%, SD ± 19.00), spending 69.92% of scans at or above the 70% synchrony threshold (including the 90% and 100% data). Cattle spent 40.43% of scans synchronized at or over the 90% threshold level, and were 100% in synchrony for 26.95% of scans. The Fleiss Kappa coefficient of agreement for 17 cattle with 33 observations and 4 categories of behavior was 0.56, indicating moderate to substantial synchrony in cattle activity. Rank affected the proportion of scans that an animal spent foraging, with lower-ranking animals spending more time foraging than higher-ranking animals (Fig. 2: GLM: LRT = 5.00, *P* = 0.03). Sex had no effect on the proportion of scans spent foraging (GLM: LRT = 0.41, *P* = 0.52).

**Fig. 2:**
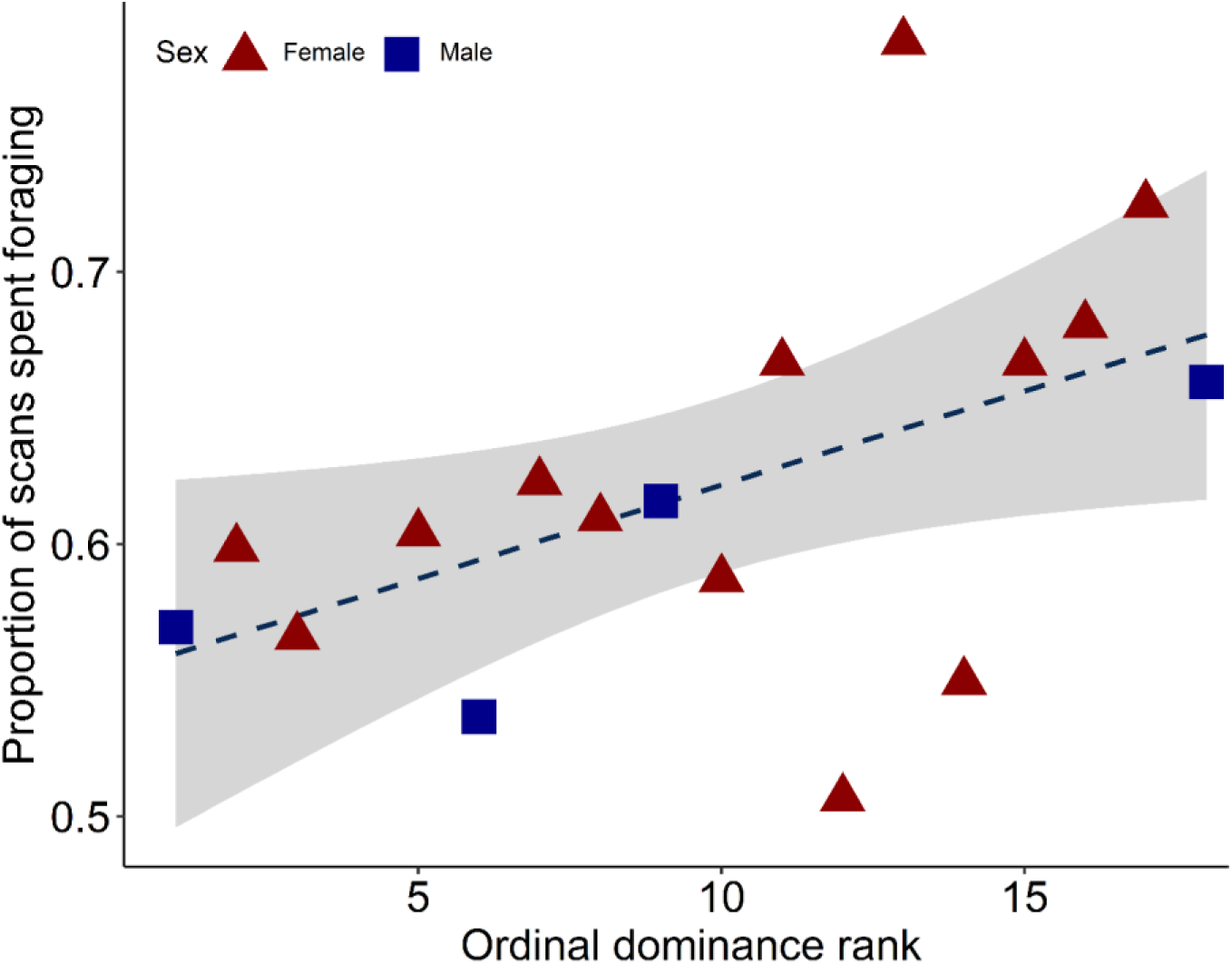
Individual proportion of scans spent foraging, relative to dominance rank, where a rank of 1 represents the most dominant animal in the group. Female raw data points are represented by red triangles, male datapoints by blue squares. Trend line is from the linear model, grey areas represent 95% confidence intervals.

### Effect of spatial proximity on synchrony (ii)

Focal animals were more synchronized with their nearest neighbors than with a random conspecific (GLMM: LRT = 68.17, *P* < 0.001). Synchrony averages between the focal animal and neighbor 1 (0.84, SD ± 0.24), neighbor two (0.80, SD ± 0.26) and neighbor 3 (0.75, SD ± 0.28) were higher than between the focal animal and the random animal (0.65, SD ± 0.32; Fig. 3).

**Fig. 3:**
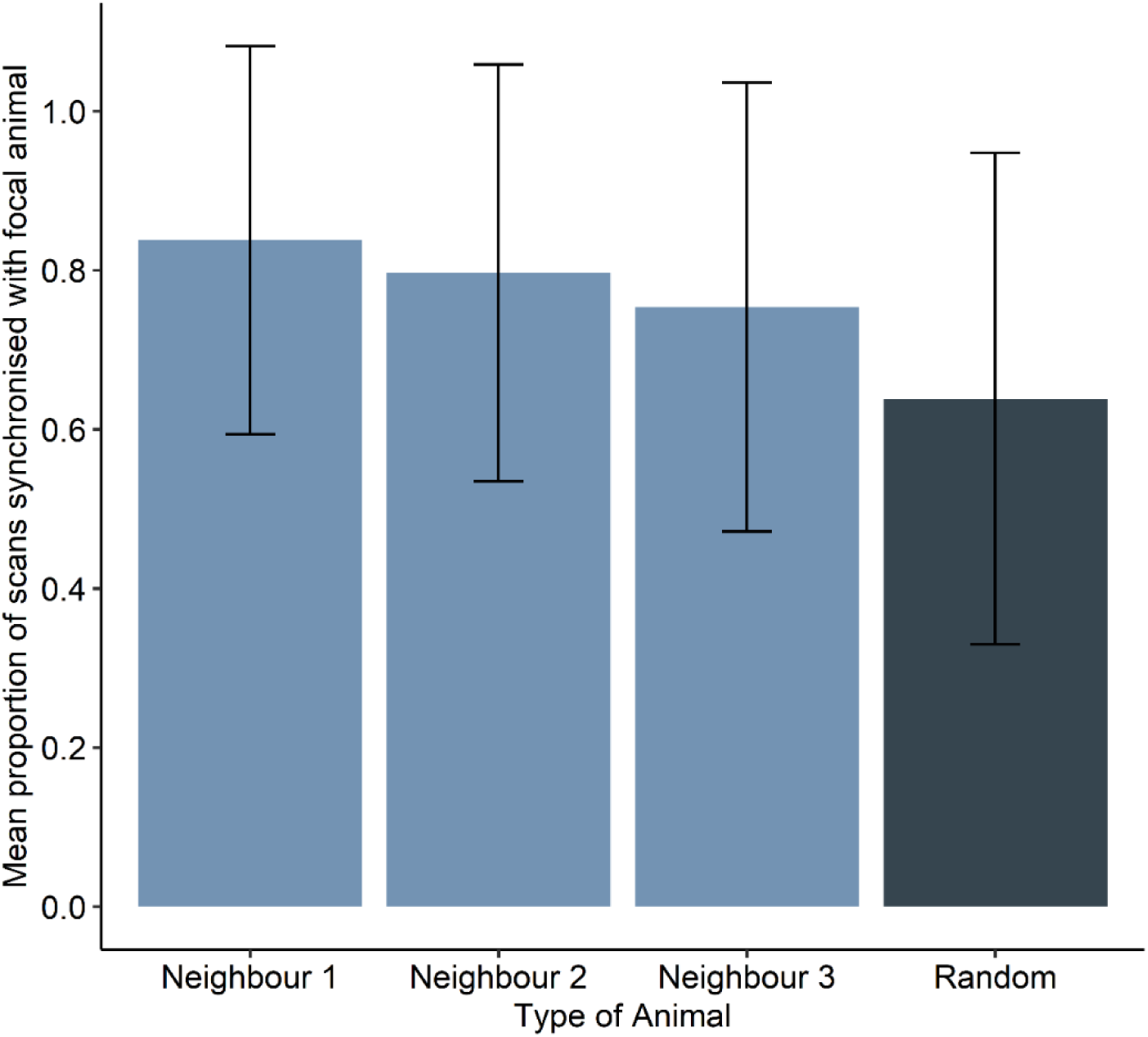
Mean proportion of scans synchronized with the focal individual per type of animal, from nearest neighbor 1 to 3, and a random conspecific. Error bars are the raw data standard deviation per group.

### Effect of sex, dominance and affiliation on synchrony and proximity (iii)

Same-sex dyads were more synchronized than different-sex dyads (LMM: *F*_1, 123_ = 6.17, *P* = 0.01), with female-female and male-male dyads having a higher proportion of synchronization than female-male dyads. There was a trend towards affiliation affecting dyadic synchrony (LMM: *F*_1, 114_ = 3.19, *P* = 0.08), but dominance rank difference did not affect dyadic synchrony (LMM: *F*_1, 114_ = 2.66, *P* = 0.11). The mean observed proportion of dyadic synchrony was 67.98%, with an expected mean dyadic synchrony of 43.00%, and dyads were more synchronized than the expected distribution of dyadic synchrony if animals were behaving independently (paired-samples t-test: t_135_ = 55.72, *P* < 0.001). The likelihood of a dyad being in close proximity (within 3 body lengths) was not related to the sex combination of the dyad (LMM: *F*_1, 118_ = 0.20, *P* = 0.66), affiliation (LMM: *F*_1, 130_ = 0.14, *P* = 0.71), or dominance rank difference (LMM: *F*_1, 131_ = 0.02, *P* = 0.90). Likewise, the synchrony between a focal animal and its nearest neighbor was not affected by previous affiliation (GLMM: LRT = 0.49, *P* = 0.48), dyadic sex combination (GLMM: LRT = 0.91, *P* = 0.34) or dominance rank difference (GLMM: LRT = 0.13, *P* = 0.72).

### Effect of synchrony on likelihood of group fission (iv)

Group synchrony was related to the likelihood of a change in group size (fission) from one scan to another (GLMM: LRT = 9.11, *P* < 0.01), with changes in group size being more likely when group synchrony was lower. Sex ratio of the scan did not affect the likelihood of group size change (GLMM: LRT = 2.31, P = 0.13).

## DISCUSSION

### Synchronization and fission dynamics

Although behavioral synchronization is vital for movement coordination and collective behavior, it is difficult to determine the relationships between proximity, individual characteristics and synchrony in wild ungulate populations (Amichay et al., 2024; Duranton & Gaunet, 2016). Feral cattle with known social relationships allow us to determine how dyadic affiliation and dominance affect the likelihood of synchrony and proximity. Our results demonstrate that sex affects the likelihood of synchrony in feral cattle, with same-sex dyads having higher synchrony than different-sex dyads. Dyads were more synchronized than expected by chance, and proximity affected the likelihood of synchrony. Surprisingly we did not find any effect of affiliation or dominance rank difference on the likelihood of a dyad being in close proximity or synchrony between nearest neighbors. Although we found that higher-ranking animals spent less time foraging, sex did not affect foraging activity. Synchronization decision-making may come with varying costs to the individual, and we find that synchrony was related to changes in group size, with fission of group members being more likely in lower synchrony scans. Our results help to elucidate the drivers of sexual segregation evolution and highlight how underlying processes can affect collective movement in group-living mixed-sex ungulates (Bonenfant et al., 2004; Ruckstuhl & Neuhaus, 2002).

### Rank and sex affect foraging activity and dyadic synchrony

In accordance with our prediction, subordinate animals spent more time foraging than dominant animals, but there was surprisingly no difference in time spent foraging between males and females. Lower-ranking animals are expected to consume lower-energy resources when higher-ranking animals are present, requiring them to spend more time foraging to compensate (Rands et al., 2006). Dominant feral cattle may monopolize resources (as found similarly in a range of social taxa, from vulturine guineafowl (*Acryllium vulturinum*; Papageorgiou and Farine 2020) to chimpanzees (*Pan troglodytes*; O’Malley et al. 2016), and disturb subordinate animals when foraging, with subordinates being more likely to leave patches of food before they are finished (Ward & Webster, 2016). Feral cattle in Hong Kong have linear, steep and stable hierarchies (Hodgson et al., 2025a), with our results indicating that dominant feral cattle can reliably use their status to monopolize resources and thus obtain fitness-related benefits from their hierarchy position. Contrary to our predictions, sex did not affect foraging activity. Although feral cattle in Hong Kong are sexually dimorphic in body size and males are larger than females, this dimorphism is moderate, and unknown factors such as health status or age may also obscure sex differences (Perroux et al., 2025b). The effect of individual traits on food access is expected to be context-dependent (Kidjo et al., 2016), and sex differences in this group may also have been obscured by the sex ratio (4 males to 13 females); further investigation into mixed-sex herds with varying sex ratios could help elucidate any effect of sex on foraging strategy. Larger rumens in male ungulates are expected to allow them to consume abundant lower-quality forage, with smaller-bodied females having larger requirements for high-quality forage, especially during reproduction (Barboza & Bowyer, 2000; Ruckstuhl, 1998). It is difficult to determine whether male and female cattle ate similar quality food items as we did not record the types of vegetation that individuals were consuming. However, individuals were observed eating similar types of vegetation, and it is unlikely animals were selecting different plants within the same area.

We found that same-sex dyads were more synchronized than animals of different sexes, highlighting how synchronization helps to shape the evolution of sexual segregation. Same-sex synchronization has been found in other ungulates (e.g., sheep, *Ovis aries*; Michelena et al. 2006), with red deer being more synchronized in single-sex groups than mixed-sex groups (Bonenfant et al., 2004). As we did not find any sex differences in foraging frequency, this indicates that segregation may occur temporally in the probability of movement, with male and female feral cattle switching activities to synchronize with others of the same-sex, but ultimately performing the behavior at the same rate (Ruckstuhl & Kokko, 2002; Sueur & Deneubourg, 2011). Our results indicate sex-biased social attraction between dyads, suggesting a social segregation transitional step between mixed-sex groups and full sex-segregation. It is not unusual for other ungulates to lack sexual differences in activity; for example, male and female mule deer (*Odocoileus hemionus*) are more efficient when foraging together (Bowyer & Kie, 2004). Social factors can also be a greater influence than ecological factors in driving sexual segregation (Galezo et al. 2018). Although we found no effect of affiliation or dominance on dyadic synchrony, social factors may play a larger role in other kinds of coordinated movement, such as leadership (Krueger et al., 2014; Sueur et al., 2018). Other social factors (such as male harassment, kinship and familiarity) may differentially affect dyads, with familiarity increasing synchronization in farmed cattle (Gygax et al., 2010), and shared stressful experiences leading to closer proximity in sheep (Keshavarzi et al., 2023). These dyadic relationships may mediate an animal’s response to neighbors in addition to sex, and species-specific differences could help identify different drivers of sexual segregation.

### Proximity affects behavioral synchrony

Proximity affected behavioral synchrony and cattle were more synchronized with nearest neighbors than with another random conspecific in the group. Spatial proximity is likely to affect how animals orientate themselves, and is a common factor in synchrony in fallow deer (Hoyle et al., 2021), red deer (Rands et al., 2014) and farm cattle (Stoye et al., 2012). Conspecific behavior provides information on habitat features, with foraging individuals providing environmental cues about food location or quality, and resource availability may drive activity and preferred space use (Dall et al., 2005). However, other ungulate behaviors, such as vigilance, are also affected by proximity and familiarity, with bighorn sheep (*Ovis canadensis*) being more likely to mimic vigilance in neighboring sheep which were spatially closer and more familiar (McDougall & Ruckstuhl, 2018). Paying attention to and responding to the behavior of a neighbor may be a feedback mechanism allowing the group to transmit information through behavioral contagion and stay cohesive; mimicry in activity thus leads to the maintenance of close proximity and subsequently further synchronization (Lakin et al., 2003; McDougall & Ruckstuhl, 2018).

The lack of relationship between proximity and any sociodemographic factor is unexpected, as well as our result that synchrony between nearest neighbors was not affected by sex, dominance nor affiliative relationship. Finer-scale individual differences may also affect proximity and animals may only change their behavior when close neighbors move out of sight. Dyadic proximity is often determined through shared motivation, and not related to the relationships between individuals (Asher & Collins, 2012). Instead of social facilitation, activity may be coordinated through collective responses to environmental cues. Animals may make independent decisions due to the influence of ecological stimuli, such as a high-quality foraging patch or an environment with reduced predation risk to rest in; however, due to the lack of large natural predators in Hong Kong, the latter may be unlikely in this study’s population (Asher & Collins, 2012; Dudgeon & Corlett, 1994). However, cattle foraging behavior has previously not been associated with vegetation characteristics (Gabrieli & Malkinson, 2024), with social associations being random in cattle when grazing in smaller groups (Stephenson et al., 2016). Without foraging and nutritional information, these nuances are difficult to determine in wild-living groups.

### Synchrony in relation to fission dynamics

The likelihood of feral cattle leaving the group was related to group synchrony, with fission being more likely when synchrony was lower, potentially reinforcing a feedback loop between the two. Our results suggest that a lack of consensus in decision-making can promote fission events and drive temporary subgrouping (Busia et al., 2022; Kerth, 2010). The feral cattle group was highly synchronized overall, with an average group synchrony of approximately 80%, matching similar values in sheep (Gautrais et al. 2007) and farm cattle (Stoye et al., 2012). All observations were mixed-sex, which unfortunately did not allow for us to investigate whether single-sex group synchrony differed from mixed-sex group synchrony. However, social factors (such as affiliative bonds, harassment or aggression) also affect fission-fusion dynamics (Darden & Croft, 2008; Galezo et al., 2018; Le Goff et al., 2024). Individual fission decisions are affected by whether other animals are also leaving (Ramos-Fernández & Morales, 2014), and conspecific familiarity and affiliation between leaders and other group members impacts the likelihood of group fission (King et al., 2008; Merkle et al., 2015). Our data may be limited by the assumption that a net group size change indicates that animals only left the group; however, this might not account for multiple animals leaving and a single animal joining. Fission decisions allow us to understand conflict in activity decisions and the evolution of sexual segregation in ungulates (Mooring et al., 2005), and further investigation into the identities of the animals leaving the group could test whether sociodemographic factors play a role in fission events.

In summary, our results support the evolution of sexual segregation through differences in activity synchronization. We suggest ungulate synchronization when living in mixed-sex groups is biased towards animals in close spatial proximity and those of the same-sex. Although external motivation can drive some level of synchrony in activity, social attraction as a mimetic behavior is an efficient individual-level mechanism driving group-level social cohesion in group-living animals. Our results highlight how synchrony can affect fission decisions and the importance of individual activity in collective ungulate decision making.

## ETHICAL STATEMENT

This work was approved by the Animal Research Ethics Sub-Committee of City University of Hong Kong (Internal Reference: A-0826).

## DATA AVAILABILITY

Data with code are accessible via an online OSF repository (Dataset, 2025; https://osf.io/gwpb5/?view_only=3f4b31f53452478896bb82a3b48da41d).

## ACKNOWLEDGEMENTS

We are grateful to Shirley Suet Ying Leung for her valuable help with data collection and the villagers of Kuk Po and the surrounding areas for their support.

## FUNDING

This work received funding from City University of Hong Kong (Grant Number 9610510).

## CONFLICT OF INTEREST DECLARATION

All authors declare they have no conflicts of interest.

## DECLARATION OF AI USE

We have not used AI-assisted technologies in this article’s creation.

## AUTHOR CONTRIBUTIONS

G.M.W.H: conceptualization, methodology, investigation, data curation, formal analysis, writing—original draft, writing—review and editing K.J.F: supervision, writing—review and editing T.P: writing—review and editing A.G.M: resources, supervision, writing—review and editing

## Notes

https://osf.io/gwpb5/?view_only=3f4b31f53452478896bb82a3b48da41d

### Competing Interest Statement

The authors have declared no competing interest.

